# RNA-deficient TDP-43 causes loss of free nuclear TDP-43 by sequestration

**DOI:** 10.1101/2022.09.06.506721

**Authors:** Sean S. Keating, Adekunle T. Bademosi, Rebecca San Gil, Adam K. Walker

## Abstract

Dysfunction and aggregation of the RNA-binding protein, TDP-43, is the unifying hallmark of amyotrophic lateral sclerosis (ALS) and frontotemporal dementia (FTD). Mechanisms and relative contributions of concurrent TDP-43 nuclear depletion, cytoplasmic accumulation, and post-translational modification to neurodegeneration remain unresolved. We employed CRISPR/Cas9-mediated fluorescent tagging to investigate how disease-associated stressors and pathological TDP-43 alter abundance, localisation, self-assembly, aggregation, solubility, and mobility dynamics of endogenous TDP-43 over time. Oxidative stress stimulated TDP-43 liquid-liquid phase separation into droplets or spherical shell-like ‘anisosomes’, which were not formed by over-expressed wild-type TDP-43. Further, nuclear RNA-binding-ablated or acetylation-mimicking TDP-43 rapidly formed anisosomes and inclusions that readily sequestered and depleted free normal nuclear TDP-43. The majority of total endogenous TDP-43 was sequestered into anisosomes, but retained high protein mobility and solubility. However, cytoplasmic RNA-deficient TDP-43 formed large phosphorylated inclusions that occasionally sequestered endogenous TDP-43, rendering it insoluble and immobile, indicating irreversible pathological transition. These findings suggest that post-translational modification and RNA-binding deficiency exacerbate TDP-43 aggregation and dysfunction by driving sequestration, mislocalisation, and depletion of normal nuclear TDP-43 in ALS and FTD.

## Introduction

Pathological aggregation of TAR DNA-binding protein 43 (TDP-43) in neurons is a key hallmark of most amyotrophic lateral sclerosis (ALS) and approximately half of frontotemporal dementia (FTD) cases (Ling et al., 2013; Neumann et al., 2006). TDP-43 is an essential nuclear DNA-/RNA-binding protein that physiologically regulates stability, transport, splicing, and translation of many thousands of mRNA transcripts (Ayala et al., 2011b; Buratti and Baralle, 2010; Polymenidou et al., 2011; Tollervey et al., 2011). In disease, TDP-43 pathology likely forms via progressive pathogenic changes to TDP-43, critically involving its depletion from the nucleus, mislocalisation, misfolding, aberrant liquid-liquid phase separation, self-assembly, post-translational modification, and terminal deposition into inclusions (Keating et al., 2022). Cytoplasmic accumulation of TDP-43 has been regarded a primary driver of toxicity and cellular dysfunction (Barmada et al., 2010; Dyer et al., 2021), however loss of normal nuclear TDP-43 has also been reported in absence of cytoplasmic TDP-43 inclusions, within post-mortem FTD (Nana et al., 2019; Vatsavayai et al., 2016), and ALS brains (Braak and Del Tredici, 2018). Indeed, TDP-43 nuclear depletion can independently mediate neurodegeneration (Igaz et al., 2011), likely via dysregulation of TDP-43-dependent transcription (Liu et al., 2019) and mRNA splicing functions (Jeong et al., 2017; Polymenidou et al., 2011; Wu et al., 2019). However, nuclear loss and cytoplasmic accumulation of TDP-43 largely occur concurrently in disease models (Igaz et al., 2011; Walker et al., 2015), such that the relative contributions of these processes to neuronal dysfunction or toxicity remain elusive. It is unclear whether upstream disease-associated stressors or pathological accumulation of post-translationally modified forms of TDP-43 drive aberrant loss, mislocalisation, or aggregation of functional endogenous nuclear TDP-43, which could explain the rapid progression of pathology formation and neurodegeneration in ALS and FTD.

Post-translational modifications (PTMs) are a major feature of TDP-43 pathology in ALS and FTD, and likely play diverse roles in TDP-43 dysfunction, aggregation, and toxicity. For example, phosphorylation of TDP-43 is a widely-recognised hallmark of disease (Neumann et al., 2006). Aberrantly acetylated TDP-43 also accumulates within cytoplasmic inclusions in ALS spinal cord tissue (Cohen et al., 2015; Kametani et al., 2016; Wang et al., 2017). Mimicking acetylation of TDP-43 via lysine-to-glutamine amino acid substitution at residues 145 and/or 192 increases formation of disease-reminiscent cytoplasmic inclusions that are phosphoTDP-43-, ubiquitin-, and Hsp70-positive (Chen and Cohen, 2019; Cohen et al., 2015; Wang et al., 2017). The location of these acetylated residues within the RNA-recognition motifs (RRMs) suggests that TDP-43 acetylation may lead to RNA-binding dysfunction (Cohen et al., 2015). Indeed, TDP-43 acetylation may decrease affinity for RNA by modifying electrostatic charges of nucleic acid-binding residues, indicated by loss of normal RNA splicing of TDP-43 target transcripts in cells expressing acetylation-mimicking TDP-43 (Cohen et al., 2015; Wang et al., 2017).

Loss of TDP-43 RNA-binding capacity, through RRM mutation or acetylation, thereby promotes aberrant self-assembly and pathological aggregation (Chen et al., 2019; Gasset-Rosa et al., 2019; Hallegger et al., 2021; Maharana et al., 2018; Mann et al., 2019; McGurk et al., 2018; Pérez-Berlanga et al., 2022; Yu et al., 2021). This is further supported by findings that TDP-43 species lacking RRMs, or containing ALS-associated RRM mutations, demonstrate high aggregation propensity and toxicity (Shenoy et al., 2020; Wei et al., 2017). Disruption of regular TDP-43-RNA interactions has revealed an intrinsic instability of the RRMs (Flores et al., 2019; Vivoli-Vega et al., 2020; Zacco et al., 2018), which influences TDP-43 folding states and phase transitions (Huang et al., 2013; Mann et al., 2019). TDP-43 RNA-binding status therefore directs self-assembly into various heterogenous biomolecular condensates, characterised by ‘liquid-like’ mobility of comprising proteins and exchange with surrounding intracellular TDP-43, mediated by liquid-liquid phase separation (LLPS) (Hallegger et al., 2021; Li et al., 2018; Molliex et al., 2015). RNA-binding-ablated or acetylation-mimicking TDP-43 variants in the nucleus have been shown to undergo a distinct form of LLPS, assembling RNA-deficient spherical shell-like structures, termed ‘anisosomes’, which require recruitment of molecular chaperones to maintain structure and solubility (Yu et al., 2021). Anisosome formation with normal endogenous TDP-43 may also be induced by proteolytic stress (Yu et al., 2021). However, real-time endogenous TDP-43 anisosome formation has not been thoroughly characterised in live cells, and it remains unclear whether other disease-associated stressors trigger anisosome assembly in human disease.

Oxidative stress is another ALS and FTD pathomechanism linking LLPS to pathological aggregation and nuclear depletion of TDP-43 (Gasset-Rosa et al., 2019; Gu et al., 2021; Maharana et al., 2018; Mann et al., 2019; McGurk et al., 2018). Oxidative stress is observed in brain, spinal cord, muscle and serum of ALS patients (McGurk et al., 2018; Ratti et al., 2020), and has been shown to induce redistribution or loss of nuclear TDP-43 as well as inclusion formation in various disease models (Ayala et al., 2011a; Dewey et al., 2011; Goh et al., 2018; Iguchi et al., 2012; Lei et al., 2018; Wang et al., 2016; Zuo et al., 2021). For example, under oxidative stress, cells form stress granules (SGs) – reversible cytoplasmic ribonucleoprotein complexes that limit non-essential protein translation by sequestering mRNAs and disassembled polyribosomes – involving recruitment of TDP-43 and other RNA-binding proteins (Colombrita et al., 2009; Fang et al., 2019; Khalfallah et al., 2018; Liu-Yesucevitz et al., 2010; Mazan-Mamczarz et al., 2006; Ratti et al., 2020). Assembly of TDP-43-containing SGs or other membraneless organelles is mediated by LLPS, which critically allows for disassembly following stress recovery (Mann et al., 2019; Zhang et al., 2019). TDP-43 may also undergo SG-independent LLPS under oxidative stress to form nuclear or cytoplasmic liquid-like droplets that do not associate with RNA substrates or SG markers (Gasset-Rosa et al., 2019; Gu et al., 2021; Mann et al., 2019). Mechanisms and interacting components for SG-independent LLPS are less clear, however may be related to stress-induced mislocalisation of RNA transcripts, as shown in induced pluripotent stem cell-derived motor neuron progenitors (Markmiller et al., 2021), which likely drives TDP-43 instability and self-assembly.

Chronic assembly of both TDP-43-positive SGs (McGurk et al., 2018; Ratti et al., 2020; Zhang et al., 2019), and RNA-/SG-independent TDP-43 structures (Chen and Cohen, 2019; Gasset-Rosa et al., 2019; Mann et al., 2019; McGurk et al., 2018; Yu et al., 2021; Zhang et al., 2019) results in conversion of reversible dynamic liquid-like structures into gel-like or solid immobile assemblies, which may potentially seed TDP-43 aggregation. Aberrant LLPS or disruption of protein dynamics may therefore lead to irreversible pathological transitions, denoted by loss of protein mobility. Interestingly, TDP-43 acetylation may be promoted by oxidative stress, and acetylation-mimicking TDP-43 forms cytoplasmic inclusions lacking SG markers (Chen and Cohen, 2019), which may be related to aberrant SG-independent LLPS. Therefore, acetylation or oxidative stress may drive TDP-43 condensation and deplete soluble free nuclear TDP-43 levels, however it is unclear whether TDP-43 LLPS and confinement within these liquid-like structures significantly alters normal TDP-43 dynamics, functional capacity, and pathological aggregation.

Depletion of functional, nuclear TDP-43 in disease could be driven by accumulation of TDP-43 protein through changes in endogenous TDP-43 expression, or sequestration into pathological inclusions or other phase-separated TDP-43 assemblies. Auto-regulation of *TARDBP* expression is mediated by TDP-43 binding its own mRNA (Ayala et al., 2011b; Hallegger et al., 2021; Koehler et al., 2022), as seen in disease models with decreased endogenous TDP-43 protein upon over-expression of wild-type or cytoplasmic TDP-43 (Igaz et al., 2011; Porta et al., 2018; Walker et al., 2015; Watkins et al., 2021; Winton et al., 2008). Additionally, it has been shown that exogenous wild-type TDP-43 becomes incorporated into cytoplasmic TDP-43 structures in cells (Che et al., 2015; Gasset-Rosa et al., 2019; Lu et al., 2021; Porta et al., 2018), resembling pathological inclusions observed in ALS and FTLD-TDP that contain natively-folded TDP-43 (Chen et al., 2019; Cohen et al., 2015; Flores et al., 2019; Maurel et al., 2018; Zacco et al., 2018). However, it is unclear whether normal endogenous TDP-43 is sequestered into pathological TDP-43 structures, and whether such sequestration could deplete overall nuclear levels and impair the dynamic properties of endogenous TDP-43.

In this study, we therefore sought to determine how disease-related stressors and RNA-binding ablation or acetylation of pathological TDP-43 affect endogenous TDP-43 levels, self-assembly, recruitment to puncta and inclusions, and mobility dynamics in the nucleus and cytoplasm of cells. We used CRISPR/Cas9 gene-editing to develop cell lines with differential fluorescent protein insertions across multiple *TARDBP* alleles. We found that oxidative stress caused fluorescently-tagged endogenous TDP-43 protein to assemble into various biomolecular condensates, including SGs, dynamic liquid-like nuclear droplets, or anisosomes, over time. Notably, over-expression and aggregation of RNA-binding-ablated, or acetyl-mimicking TDP-43 drives nuclear depletion of endogenous TDP-43, via concentration-dependent sequestration into inclusions, puncta or anisosomes. This specific sequestration into large, phosphorylated nuclear or cytoplasmic inclusions formed by acetylated TDP-43 led to insolubility and immobility of endogenous TDP-43, indicating irreversible pathological transition. The most dramatic sequestration was caused through formation of nuclear anisosomes comprised of acetylation-mimicking TDP-43, while endogenous TDP-43 recruited to these dynamic structures remained highly mobile. Our findings therefore suggest that both acetylation and RNA-binding ablation increase TDP-43 aggregation propensity, and also cause a loss of free normal nuclear TDP-43 by promoting its mislocalisation and sequestration into pathological assemblies, thereby driving TDP-43 dysfunction and further aggregation in ALS and FTD.

## Results

### Generation and validation of TDP-43-reporter cell lines using CRISPR/Cas9-mediated multiple allele fluorescence tagging of the endogenous *TARDBP* locus

To visualise and quantify the spatiotemporal dynamics of endogenous TDP-43, we inserted fluorescent tags to the *TARDBP* C-terminus in single or multiple alleles using CRISPR/Cas9-mediated homology-directed repair (Figure 1A). We exploited the hypertriploid karyotype (i.e., multiple copies of the *TARDBP* allele) of HEK293T cells (Lin et al., 2014) to generate three different fluorescently-tagged *TARDBP* cell lines, by mixing *mAvicFP1* and *mCherry* DNA donor templates in a single reaction (Figure 1B,C; Supplementary Figure 1). mAvicFP1 is a recently discovered fluorescent protein, derived from *Aequorea victoria*, which has 80% homology to traditional GFP, but produces brighter chromophores and is amenable to single-molecule microscopy (Lambert et al., 2019). Our approach resulted in a polyclonal population of cells with expression of TDP-43 tagged with mAvicFP1 or mCherry, either alone or in combination, from donor integration across multiple *TARDBP* alleles (Figure 1D). Fluorescence-activated cell sorting revealed 4.74% of cells were mAvicFP1-positive, 4.17% were mCherry-positive, and 0.54% were both mAvicFP1- and mCherry-positive, or ‘dual-tagged’, with the remainder (90.5%) unedited (Figure 1E).

**Figure 1.**
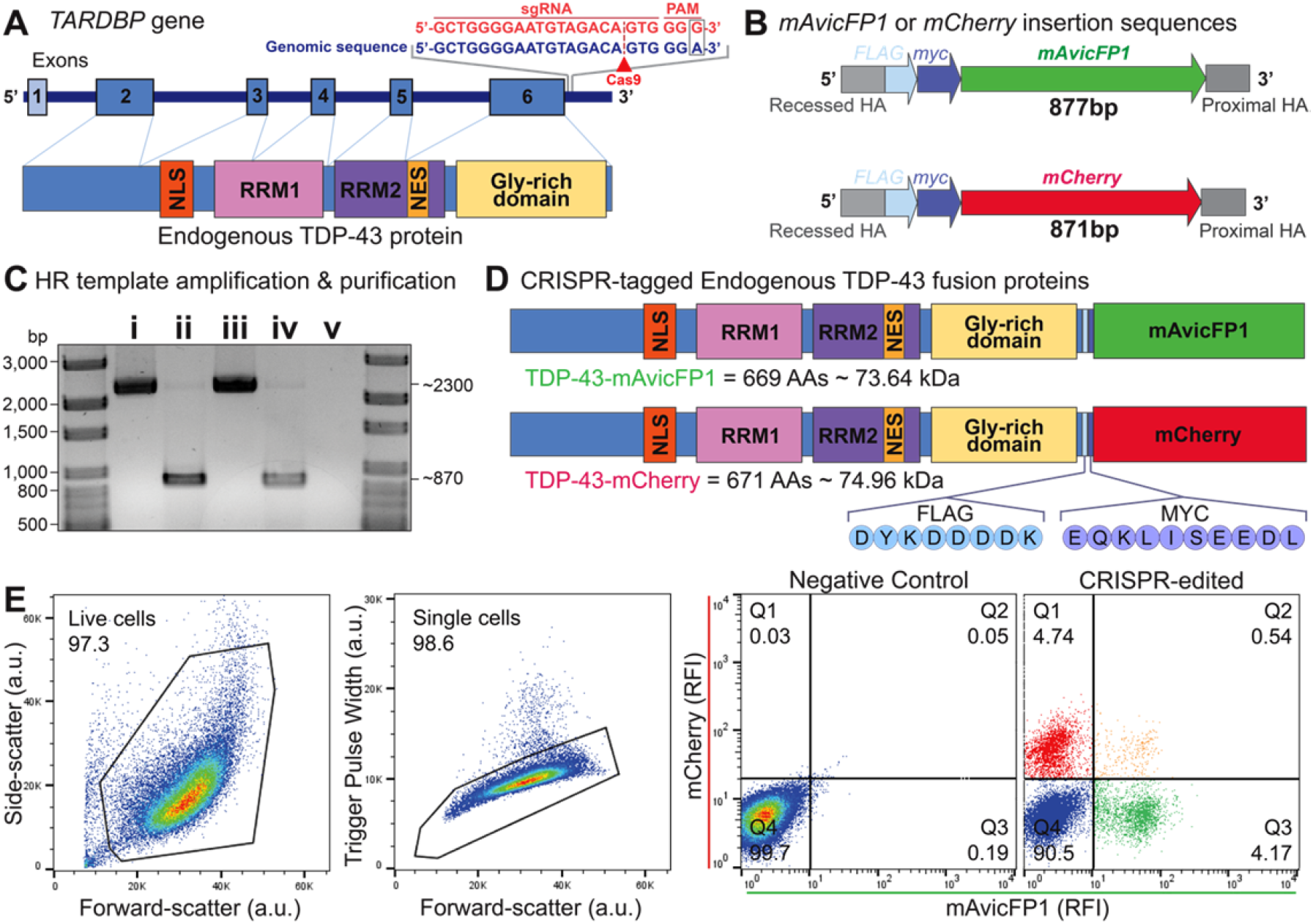
CRISPR/Cas9-mediated gene editing for single and dual insertion of *mAvicFP1* or *mCherry* to endogenous *TARDBP* C-termini. **(A)** Schematic of endogenous *TARDBP* gene and protein structure, including the sgRNA target site and mutated PAM sequence (red text) that guides Cas9 (red arrowhead) for DNA double-stranded breaks at the C-terminus of *TARDBP* (blue text). **(B)** Schematic of double-stranded DNA donor insertion sequence design and **(C)** DNA electrophoresis of **(i)** mAvicFP1 plasmid, **(ii)** mAvicFP1 PCR amplicon donor template, **(iii)** mCherry plasmid, **(iv)** mCherry PCR amplicon donor template, and **(v)** non-template control. **(D)** Schematic of the expected protein products of CRISPR/Cas9-mediated gene insertion, endogenous TDP-43-mAvicFP1 and TDP-43-mCherry fusion proteins, including amino acid (AA) length and predicted protein size. **(E)** Fluorescence-activated cell sorting of nucleoporated HEK293T cells. *Left to right*: plot of forward- and side-scatter resolves live cells (a.u., arbitrary units), plot of forward-scatter and trigger pulse width resolves single cells. Representative plots of mAvicFP1 and mCherry relative fluorescence intensity (RFI) from negative control and polyclonal CRISPR-edited samples. Quadrant gating was set using the negative control.

To validate the efficiency and accuracy of the multi-allelic CRISPR/Cas9 gene insertion, we analysed RIPA-soluble protein and genomic DNA from two candidate clonal cell lines that were of low-, medium-, or high-fluorescence intensity for each of *TARDBP*-*mAvicFP1, TARDBP*-*mCherry*, or dual-tagged *TARDBP*-*mAvicFP1* plus *TARDBP*-*mCherry*. Immunoblotting for TDP-43 revealed multiple bands that differed in molecular weight, including ‘untagged’ TDP-43 (43 kDa) or high-molecular-weight tagged TDP-43-mAvicFP1 (∼72kDa) and TDP-43-mCherry (∼74 kDa) fusion proteins (Figure 2A). Probing for myc, which was inserted in the linker sequence, confirmed that higher-molecular-weight TDP-43 bands were CRISPR edited fluorescently-tagged fusion proteins. To estimate the proportion of *TARDBP* loci with or without insertion, we calculated an ‘editing ratio’ for each clone, by dividing the combined densitometry signals of 72/74 kDa tagged-TDP-43 by the sum of all TDP-43 bands (Figure 2B-D, Supplementary Figure 2). For example, an editing ratio of 1.0 indicates complete CRISPR insertion across all alleles, as seen with *TARDBP-mAvicFP1/-mCherry* clone C4 (Figure 2D), which also showed the highest fluorescence intensity of the representative clones (Figure 2E). *TARDBP*-mCherry clone B8 had an editing ratio of ∼0.5, suggesting that half of *TARDBP* alleles possessed an *mCherry* tag (Figure 2A and B). *TARDBP* editing ratio generally correlated with fluorescence intensity of each monoclonal cell line observed by confocal microscopy, which also revealed the expected sub-nuclear distribution of TDP-43 with exclusion from nucleoli (Figure 2E). Finally, we performed Sanger sequencing on genomic DNA from clonal cell lines of the highest editing efficiency. The DNA donor templates, including *FLAG-myc* linker and *mAvicFP1* or *mCherry* sequences, were precisely inserted at the intended C-terminal *TARDBP* Exon 6 locus (Supplementary Figure 3).

**Figure 2.**
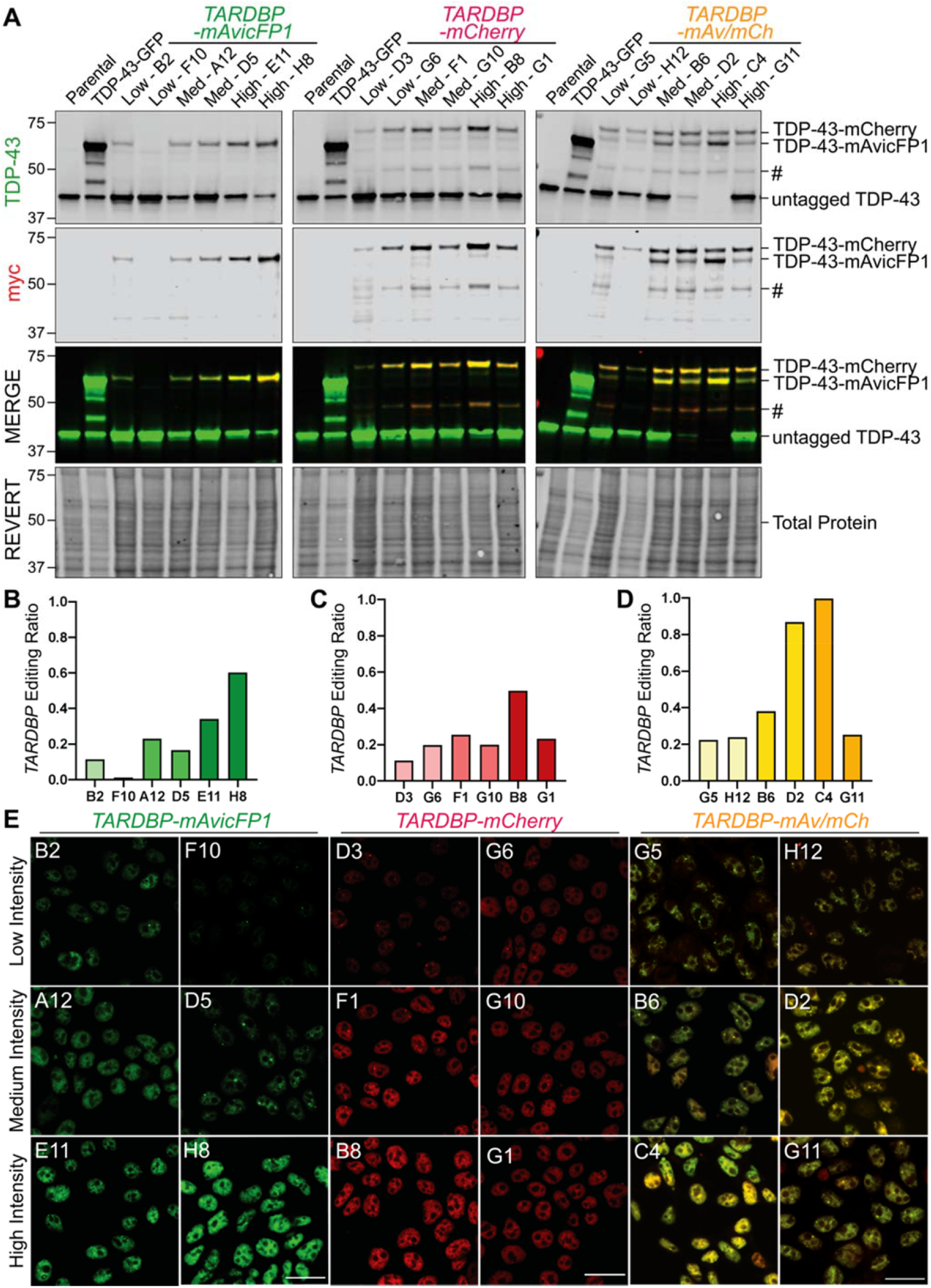
Expression of high-molecular weight TDP-43 protein verifies successful CRISPR/Cas9-mediated gene insertion of *mAvicFP1* and/or *mCherry* to *TARDBP* alleles. Two monoclonal cell lines of low, medium, or high intensity were selected. **(A)** Immunoblotting for TDP-43 and myc revealed the abundance of untagged (43kDa) and tagged TDP-43 (72-74kDa), which was quantified by densitometry analysis. CRISPR-editing ratios for **(B)** *TARDBP-mAvicFP1*, **(C)** *TARDBP-mCherry*, or **(D)** *TARDBP-mAvicFP1/-mCherry* monoclonal cell lines were calculated by dividing the total protein-normalised signals of high-molecular weight tagged TDP-43 by the combined signals of all TDP-43 bands, as described in Supplementary Figure 2. # represents a lower-molecular-weight band positive for endogenous mCherry-tagged TDP-43. **(E)** Representative confocal images captured at 63x magnification depicting relative fluorescence intensities of *TARDBP*-tagged cell lines, with clone names indicated in the upper left corner. Note that *TARDBP-mAv/mCh* images are displayed as a merge of separate mAvicFP1 and mCherry channels; scale = 20μm.

**Figure 3.**
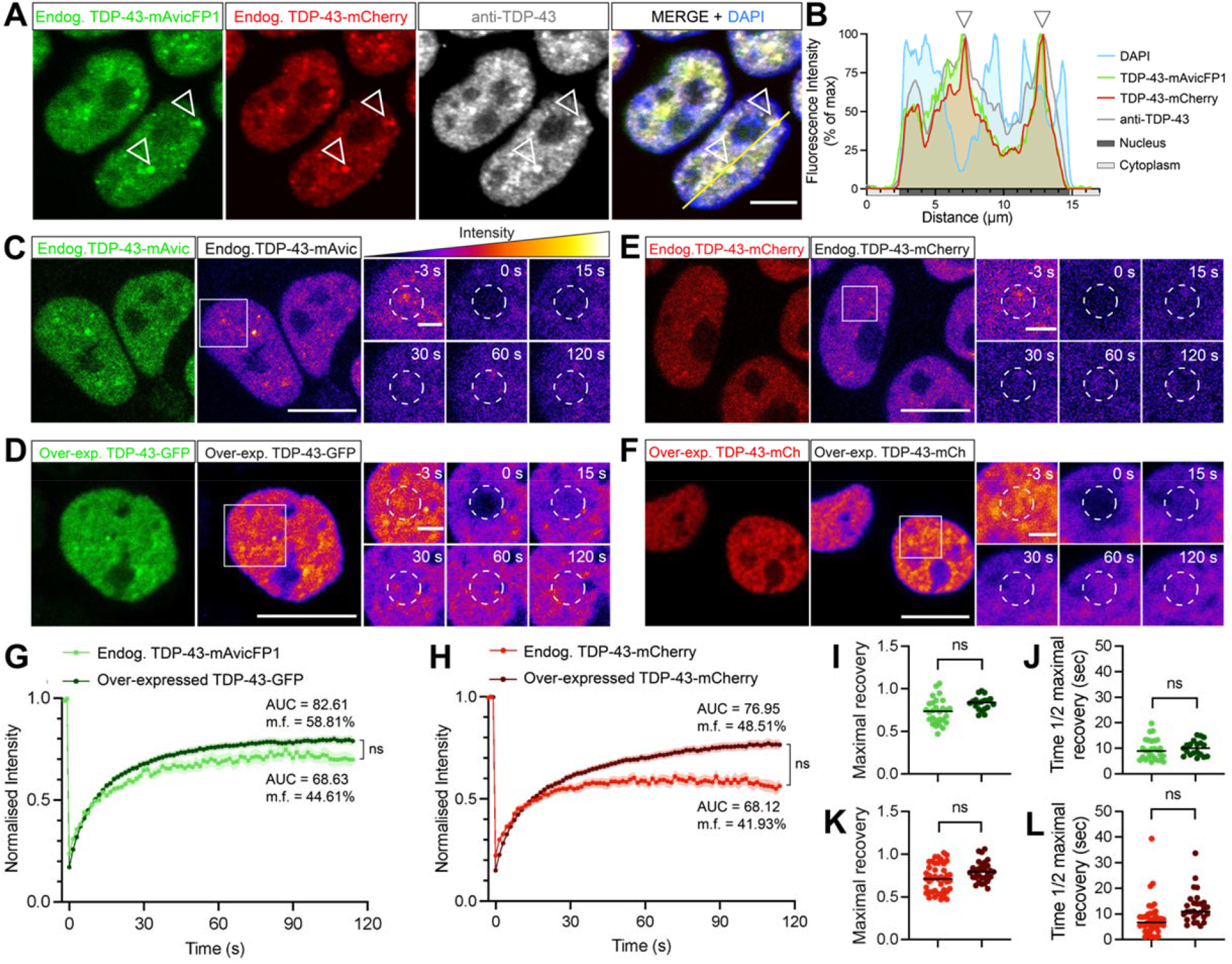
CRISPR-tagged endogenous TDP-43-mAvicFP1/-mCherry fusion proteins demonstrate nuclear localisation and high protein mobility under physiological conditions. **(A)** CRISPR-tagged TDP-43-mAvicFP1 and TDP-43-mCherry proteins exhibit nuclear localisation and co-localise strongly with anti-TDP-43 labelled TDP-43, indicated by arrowheads. Scale bar = 5μm. **(B)** Intensity profile displaying spatial alignment of TDP-43-mAvicFP1, TDP-43-mCherry and anti-TDP-43 immunofluorescence signals within nuclei. **(C-F)** FRAP was performed to quantify the mobility of nuclear endogenous TDP-43 in *TARDBP-mAvicFP1* cells or *TARDBP-mCherry* cells. The two large left panels (scale bar = 10μm) show initial snapshots of **(C)** endogenous TDP-43-mAvicFP1, **(D)** over-expressed TDP-43-GFP, **(E)** endogenous TDP-43-mCherry, and **(F)** over-expressed TDP-43-mCherry, depicted with the correct emission wavelength (green or red) on the left and pseudocoloured “fire” LUT to visualize the scale of intensity on the right. Image intensities were re-scaled between conditions to allow for side-by-side comparison of fluorescence recovery. Smaller panels display a zoomed-in view of the bleached region of interest outlined by white dashed circles, captured at -3, 0, 15, 30, 60 and 120 s post-bleaching (scale bar = 2μm). **(G, H)** Normalised intensity over time post-bleaching, with mobility fraction (m.f.) values and area-under-the-curve (AUC) averaged for each cell, which was used for statistical comparison. Data is the mean of *n*=3 biologically independent repeats, each including at least 5 cells on average, and the shaded area denotes standard error about the mean. **(I, K)** Maximal fluorescence recovery at 120 s post-bleaching, for mAvicFP1/GFP and mCherry, respectively. **(J**, **L)** Time taken to recover to half of the maximal level, for mAvicFP1/GFP and mCherry, respectively. Black lines represent mean of n=3 independent repeats as above, with each dot representing individual FRAP recordings across multiple cells in each replicate. Data analysed using two-tailed t-tests where ns is not significant (p > 0.05).

### Fluorescently-tagged endogenous TDP-43 exhibits characteristic protein localisation, self-assembly, and mobility dynamics under physiological conditions or oxidative stress

To determine whether characteristic changes in TDP-43 localisation, dynamics, and aggregation under physiological conditions and disease-associated stress were recapitulated in our CRISPR-tagged endogenous TDP-43 cell lines, we performed a series of imaging assays. TDP-43 immunolabelling of *TARDBP-mAvicFP1/-mCherry* cells demonstrated clear co-localisation between the anti-TDP-43 immunostaining pattern, TDP-43-mAvicFP1, and TDP-43-mCherry fusion proteins (Figure 3A and B). Fluorescence recovery after photobleaching (FRAP) showed that diffusely distributed nuclear endogenous TDP-43-mAvicFP1 and TDP-43-mCherry exhibit similar nuclear localisation and overall mobility dynamics compared to exogenous over-expressed wild-type TDP-43-mGFP or TDP-43-mCherry (Figure 3C-L). No significant difference was found in the maximal recovery levels normalised to pre-bleaching intensity (0.74 and 0.83 for endogenous TDP-43-mAvicFP1 and over-expressed TDP-43-GFP, respectively [Figure 3I], or 0.76 and 0.80 for endogenous and over-expressed TDP-43-mCherry, respectively [Figure 3J]). Regarding recovery rate kinetics, there was no significant difference in time taken to reach half maximal recovery between endogenous and over-expressed fusion proteins (8.71 s and 9.81 s, respectively, for mAvicFP1 or GFP TDP-43 [Figure 3K]; 8.5 s and 11.49 s, respectively, for mCherry TDP-43 [Figure 3L]). Area-under-the-curve (AUC) and mobility fraction (m.f.) analyses, which further indicate the overall proportion of highly mobile proteins within the nucleus, were slightly but insignificantly decreased for both endogenous fusion proteins relative to their over-expressed forms (AUC = 67.80 and 82.61, m.f. = 44.61 and 58.81% respectively, for mAvicFP1 and GFP TDP-43 [Figure 3G]; AUC = 68.12 and 76.95, m.f. = 41.93 and 48.51% respectively, for mCherry TDP-43 [Figure 3H]). Therefore, CRISPR-tagged endogenous TDP-43 proteins demonstrate similar properties to commonly-studied over-expressed TDP-43 fusion proteins, thereby validating these cell lines for further studies of TDP-43 protein dynamics.

Liquid-liquid phase separation (LLPS) is a key feature of physiological TDP-43 function, but aberrant or chronic self-assembly via LLPS may seed aggregation of TDP-43. As TDP-43 LLPS has commonly been studied using over-expression constructs, truncated mutants, optogenetic clustering methods, or recombinant proteins *in vitro*, we sought to characterise LLPS structures comprising endogenous native TDP-43 protein in live cells. As an essential RNA-binding protein, TDP-43 has previously been shown to be a major component of SGs that assemble under oxidative stress (Lee et al., 2021; Mann et al., 2019; Ratti et al., 2020; Zhang et al., 2019). Indeed, fixed-cell immunostaining for the SG marker, G3BP1, in *TARDBP-mAvicFP1* cells under physiological conditions demonstrated absence of cytoplasmic SGs with diffusely distributed endogenous TDP-43 in the nucleus (Figure 4A). As expected, treatment with the oxidative stressor, 50 μM sodium arsenite, for 1 h induced formation of distinct rounded G3BP1-positive and endogenous TDP-43-positive puncta of ∼2-4 μm in diameter (Figure 4B). Live-cell imaging of vehicle-treated cells also revealed small nuclear physiological endogenous TDP-43 puncta, defined as rounded particles of 0.5-2.5 μm in diameter (Figure 4C). Arsenite-mediated oxidative stress increased the percentage of cells with nuclear endogenous TDP-43 puncta by approximately 20%, and the number of puncta per cell by 50% (Figure 4 D-H). However, arsenite-induced cytoplasmic endogenous TDP-43 SGs could not be detected under live-cell acquisition parameters.

**Figure 4.**
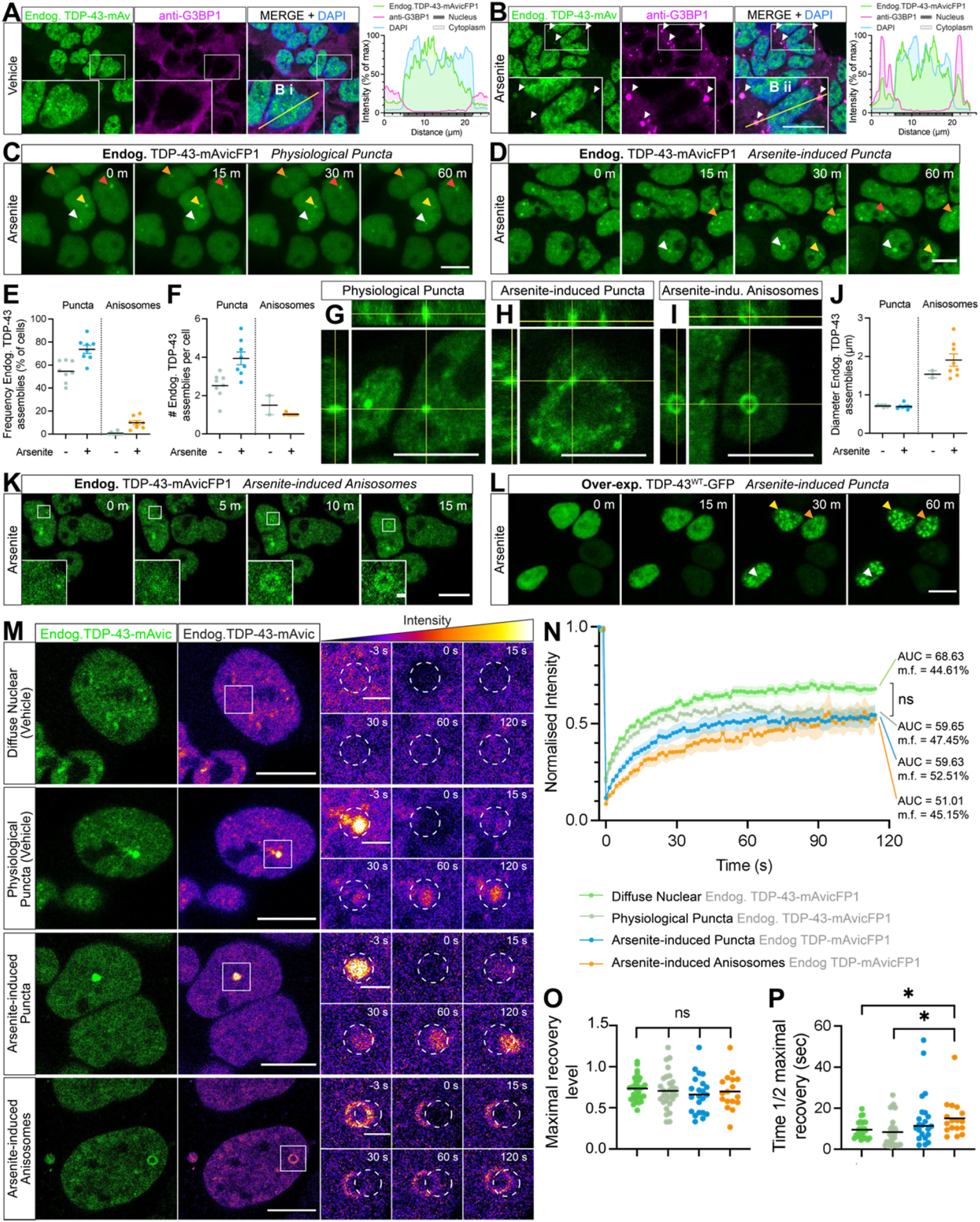
Endogenous CRISPR-tagged normal nuclear TDP-43 undergoes physiological or stress-induced self-assembly via liquid-liquid phase separation to form cytoplasmic stress granules or dynamic nuclear puncta and anisosomes in real-time. **(A, B)** Representative confocal micrographs and intensity profiles of anti-G3BP1 labelling and TDP-43-mAvicFP1 in HEK293:*TARDBP-mAvicFP1* cells treated with vehicle or 50 μM sodium arsenite for 1 h. Inset zoomed view of cells measured for fluorescence intensity profiles; scale = 10 μm. **(C)** Live-cell imaging displays formation of stable physiological puncta with 1 h vehicle treatment, or **(D)** increased formation of nuclear puncta induced by 50 μM arsenite. **(E)** Frequency of cells with endogenous TDP-43-mAvicFP1 nuclear puncta and **(F)** number of puncta per cell. Data points represent the mean of all cells within an image, totalling 8 images per group. **(G)** 3-dimensional orthogonal projections of physiological nuclear puncta **(H)** arsenite-induced puncta, and **(I)** arsenite-induced anisosomes comprising endogenous TDP-43-mAvicFP1. **(J)** Diameter size in μm of puncta and anisosomes. **(K)** Live-cell imaging of endogenous TDP-43-mAvicFP1 anisosome formation with 50 μm arsenite treatment. **(L)** Live-cell imaging of parental HEK293 cells over-expressing TDP-43WT-GFP with 50 μm arsenite treatment; scale large panels = 10 μm; zoomed panels = 1 μm. **(M)** FRAP quantification of endogenous TDP-43-mAvicFP1 mobility within distinct nuclear assemblies. Two large left panels depict endogenous TDP-43-mAvicFP1 singal with correct emission (green) on the left and pseudocoloured “fire” LUT to visualise intensity scale on the right; scale = 10 μm. Image intensities re-scaled between conditions to allow for side-by-side comparison of fluorescence recovery. Smaller panels display zoomed view with photo-bleached ROI outlined by white dashed circles, captured at -3, 0, 15, 30, 60 and 120 s post-bleaching; scale = 2 μm. **(N)** FRAP analysis displays normalised intensity during recovery over 120 s, along with mobility fraction (m.f.) values and area-under-the-curve (AUC) averaged for each cell, which was used for statistical comparison. Coloured lines represent mean of *n*=3 biologically independent repeats, each striving for at least 5 cells, with shaded areas denoting standard error about the mean. **(O)** Maximal fluorescence recovery at 120 s post-bleaching, and **(P)** time taken to recover to half of the maximal level for each of the TDP-43-mAvicFP1 assemblies. The same colour scheme as that of the intensity line graphs was used, with black lines representing the mean across n=3 biologically independent repeats, and each dot denoting individual cells. All data analysed with one-way ANOVA multiple comparisons with Tukey’s post hoc test, relative to the ‘diffuse nuclear endogenous TDP-43-mAvicFP1’ control (* = p < 0.05).

Interestingly, arsenite treatment in *TARDBP-mAvicFP1* cells also induced formation of assemblies reminiscent of the recently characterised ‘TDP-43 anisosomes’, spherical shell-like structures with a core void of TDP-43 (Figure 4I). Endogenous TDP-43 anisosomes formed in 10% of arsenite-treated cells, wherein most cells contained just one anisosome, compared to 0.5% of vehicle-treated cells exhibiting spontaneous anisosome formation (Figure 4E,F). Anisosomes were larger than puncta and treatment with sodium arsenite yielded anisosomes that were on average 2 μm in diameter (Figure 4J). Endogenous TDP-43 anisosomes formed rapidly (within 15 min, Figure 4K) and remained stable throughout arsenite treatment (Supplementary Figure 4A). TDP-43 anisosome formation and stability are thought to critically require Hsp70 recruitment, however immunostaining only revealed endogenous Hsp70 surrounding the periphery, but not within the core, of endogenous TDP-43 anisosomes (Supplementary Figure 4B), as shown previously (Yu et al., 2021).

Over-expression of wild-type TDP-43-GFP in parental HEK293 cells led to variable intensity levels and occasional mislocalisation or aggregation of TDP-43^WT^-GFP under basal conditions, while arsenite-mediated oxidative stress increased the brightness and number of TDP-43^WT^-GFP puncta per cell over time (Figure 4L, Supplementary Figure 4C). Notably, exogenous over-expressed TDP-43^WT^-GFP did not form anisosomes under arsenite treatment. This difference between endogenous and over-expressed TDP-43 self-assembly may highlight the dependency of TDP-43 self-assembly patterns on protein abundance. This suggests that the formation of stress-induced anisosomes and other dynamic structures composed of wild-type TDP-43 may be best studied in models that visualise physiological TDP-43.

We next used FRAP to determine the relative mobility dynamics of the distinct nuclear assemblies of endogenous TDP-43, namely physiological puncta, arsenite-induced puncta, and arsenite-induced anisosomes (Figure 4M). Endogenous TDP-43 within each of these assemblies exhibited dynamic exchange and/or recruitment of TDP-43 molecules over time, showing no significant differences in area-under-the-curve, or mobility fraction analyses (Figure 4N), and comparable maximal fluorescence recovery levels to diffusely-distributed nuclear TDP-43 (Figure 4O). However, endogenous TDP-43 anisosomes showed significantly decreased rate of recovery, measured as the time to reach half maximal intensity post-bleaching (t1/2 = 19.53 s), compared to diffuse and physiological puncta of endogenous TDP-43 (t1/2 = 7.96 s) (Figure 4P).

Together, these results demonstrate that CRISPR/Cas9-tagged endogenous TDP-43-mAvicFP1 and TDP-43-mCherry proteins exhibit characteristic TDP-43 localisation and mobility, however there were some key differences in the propensity for self-assembly under oxidative stress compared to over-expressed wild-type TDP-43. Notably, we showed that under oxidative stress, over-expressed TDP-43^WT^-GFP demonstrated increased nuclear aggregation compared to endogenous TDP-43-mAvicFP1, and did not form anisosomes which were observed in 10% of *TARDBP-mAvicFP1* cells. This indicates that over-expressed proteins may not always faithfully recapitulate self-association states of TDP-43, which may be influenced by protein concentration. Endogenous TDP-43 anisosomes were dynamic, yet demonstrated a slower fluorescence recovery rate than diffuse or physiological puncta of endogenous TDP-43, indicating decreased rate of recruitment. Therefore, both cellular stress conditions and the distinct structures of TDP-43 assemblies affects the real-time formation and mobility dynamics of normal nuclear TDP-43.

### Pathological accumulation of RNA-binding-ablated or acetylation-mimic TDP-43 sequesters and depletes free normal nuclear TDP-43 in a concentration-dependent manner

Another key feature of disease is the depletion of normal nuclear TDP-43, which is observed in cells that exhibit pathological aggregation, but can also independently mediate neurodegeneration. To determine whether the progressive accumulation of different pathological TDP-43 proteins causes loss of free nuclear TDP-43 in single cells, we over-expressed nuclear wild-type (WT), RNA-binding-ablated (4FL; F147L/F149L/F229L/F231L) or acetylation-mimic (2KQ; K145Q/K192Q) TDP-43 mutants, as well as cytoplasmically-mislocalised variants of each mutant, harbouring disrupted nuclear localisation signals (ΔNLS; K82A/R83A/K84A), in HEK293T:*TARDBP-mCherry* cell lines (Figure 5A and B). In duplicate experiments, parental HEK293T cells lacking CRISPR-tagged endogenous TDP-43-mCherry transfected with TDP-43-GFP variants were used as imaging controls to ensure no cross-over of fluorescence emission and for setting intensity thresholds during quantitative image analysis (Supplementary Figure 5). Confocal imaging and CellProfiler image analysis demonstrated that over-expression of TDP-43-GFP variants led to slight, but not statistically significant, decreases in total nuclear endogenous TDP-43-mCherry signal (white arrowheads; Figure 5A-C), compared to GFP-transfected control cells. Auto-regulation of TDP-43 is a major contributor to the loss of normal nuclear TDP-43 levels observed in cells with accumulation of pathological TDP-43, by decreasing endogenous *TARDBP* expression (Ayala et al., 2011b; Hallegger et al., 2021; Koehler et al., 2022). We found no significant change in endogenous *TARDBP-mCherry* mRNA expression in bulk-cell qPCR analysis following introduction of these exogenous TDP-43-GFP variants, including RNA-binding-competent nuclear WT TDP-43-GFP (Supplementary Figure 6A). However, this result is likely due to inconsistent exogenous TDP-43-GFP over-expression and consequent sampling of many untransfected cells in our experiments (Supplementary Figure 6B,C).

**Figure 5.**
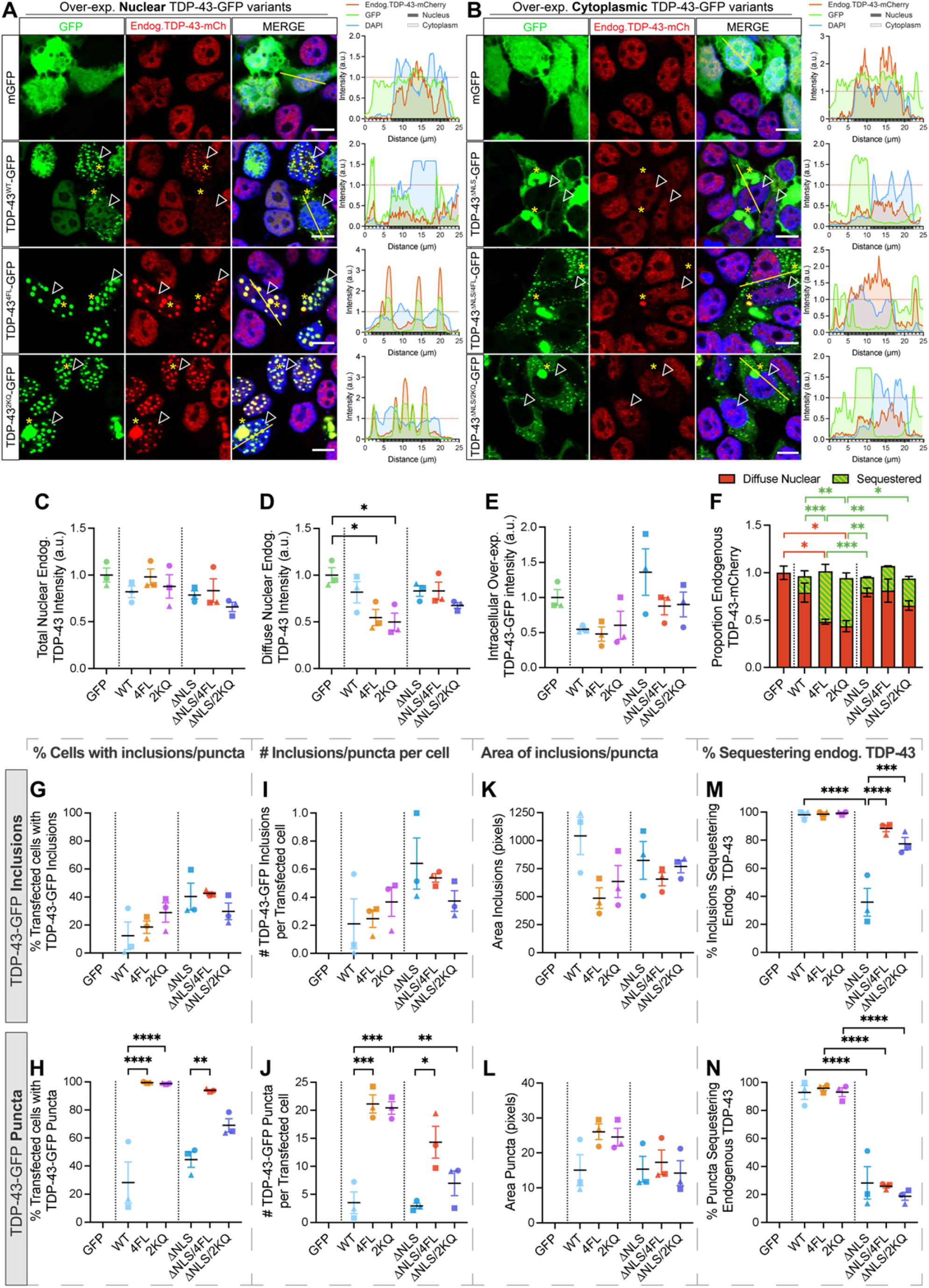
Nuclear or cytoplasmic acetylation-mimicking TDP-43 decreases endogenous TDP-43 levels and recruits endogenous TDP-43 to puncta and inclusions. Representative confocal micrographs of HEK293T:*TARDBP-mCherry* cells over-expressing **(A)** nuclear or **(B)** cytoplasmic TDP-43-GFP variants fixed and imaged at 48 h post-transfection to observe changes in levels or localisation of endogenous TDP-43-mCherry; scale = 10 μm. Right-hand panels display relative fluorescence intensity profiles of example cells (sampling region marked by yellow line), and the red line denotes the mean level of each intensity measurement across GFP-transfected control cells, to which other intensities are normalised. **(C)** CellProfiler image segmentation analysis was used to quantify intensities of total nuclear or **(D)** diffuse nuclear endogenous TDP-43-mCherry proteins, and **(E)** intracellular levels of over-expressed TDP-43-GFP protein variants in transfected cells. Each point represents the mean relative fold change from GFP-transfected control cells, sampling cells from 8 regions-of-interest at 63x magnification, for each of *n*=3 biologically independent repeats, denoted by different point shapes. **(F)** Stacked bar graph demonstrating the proportions of endogenous TDP-43 diffusely distributed in the nucleus (red) or sequestered into nuclear or cytoplasmic TDP-43-GFP aggregates (green). **(G-N)** CellProfiler image segmentation was used to quantify parameters of aggregates formed by over-expression of TDP-43-GFP variants. Inclusions were defined as bright speckles with minimum diameter of 20 pixels, and puncta with 3-20 pixel diameter, within transfected cells. **(G**,**H)** Proportion of transfected cells with inclusions or puncta. **(I**,**J)** Number of inclusions or puncta per transfected cell. **(K**,**L)** Area of inclusions or puncta. **(M**,**N)** Proportion of inclusions or puncta sequestering endogenous TDP-43, respectively. Inclusions or puncta sequestering endogenous TDP-43-mCherry defined as those with endogenous TDP-43-mCherry intensity above the lower quartile intensity of diffusely-distributed nuclear endogenous TDP-43-mCherry in GFP-transfected control cells. Cell numbers and replicates as per (C-E). Statistically significant differences between the means were analysed as one-way ANOVA multiple comparisons with Tukey’s post hoc test (* = p<0.05, ** = p<0.01, *** = p<0.001, **** = p<0.0001).

Despite no change in total levels, we observed that the distribution of endogenous TDP-43-mCherry was dramatically altered and co-localised with bright mutant TDP-43-GFP aggregates formed by each of the nuclear or cytoplasmic variants (yellow asterisks and intensity profiles; Figure 5A and B). This indicated sequestration, which was extremely pronounced in cells over-expressing nuclear RNA-binding ablated (4FL) or acetylation-mimicking (2KQ) mutants. Sequestration with 4FL or 2KQ mutants correlated with a significant 50% decrease in levels of diffuse nuclear endogenous TDP-43 (excluding areas occupied by aggregates; Figure 5D; p < 0.05). We found a clear negative correlation between GFP and diffuse nuclear mCherry intensity in individual cells with all TDP-43-GFP variants tested (Supplementary Figure 6D). This indicates that increasing intracellular levels of exogenous TDP-43-GFP resulted in a depletion of diffuse endogenous normal nuclear TDP-43 in a concentration-dependent manner (Supplementary Figure 6D). We analysed individual inclusions or puncta and measured endogenous TDP-43 intensity within these aggregates (Supplementary Figure 7). Combining the levels of ‘sequestered’ and ‘diffuse’ proportion of endogenous TDP-43 demonstrated that total cellular abundance of endogenous TDP-43 was unaffected, and that this re-distribution into inclusions or puncta accounts for the depletion of free diffuse normal nuclear endogenous TDP-43 (Figure 5F). At the single-cell level, there was a positive correlation between intensity of inclusions or puncta formed from over-expressed TDP-43-GFP and endogenous TDP-43-mCherry intensity levels sequestered into these structures (Supplementary Figure 6E and F). This shows that endogenous TDP-43-mCherry is specifically sequestered into pathological TDP-43 structures, and that nuclear RNA-binding-ablated or acetylation-mimicking TDP-43-GFP mutants exhibited dramatically increased tendencies to sequester endogenous TDP-43 (Supplementary Figure 6G and H), resulting in the greatest depletion of diffuse normal nuclear TDP-43.

All TDP-43-GFP variants demonstrated variable tendencies for forming large inclusions (Figure 5G), and the number of inclusions per transfected cell (Figure 5I), which were slightly higher for cytoplasmic mutants. However, RNA-binding-ablation (4FL) or acetylation-mimic (2KQ) variants dramatically increased formation of smaller puncta in the nucleus, which was matched only by ΔNLS/4FL, but not ΔNLS or ΔNLS/2KQ mutants in the cytoplasm (Figure 5H,J). Area of inclusions was not significantly different between variants (Figure 5K), however ‘puncta’ formed by nuclear 4FL or 2KQ mutants demonstrated larger area compared to other variants (Figure 5L) and consistently resembled anisosomes (Supplementary Figure 6I). Our results support previous work (Yu et al., 2021), showing that nuclear RNA-binding-ablated and acetylation-mimicking TDP-43 form anisosomes, which we observed in 100% of transfected cells, each containing approximately 20 anisosomes (Supplementary Figure 6J-L). Similarly, we performed Hsp70 immunostaining in these cells but were unable to verify its localisation within the core of acetylated TDP-43-GFP anisosomes that recruited endogenous TDP-43 (Supplementary Figure 6I). Inclusions and puncta sequestering endogenous TDP-43 were identified as those with TDP-43-mCherry signals above lower quartile intensity of nuclear endogenous TDP-43-mCherry in GFP-transfected control cells. Nearly all nuclear inclusions formed by WT, 4FL or 2KQ variants sequestered endogenous TDP-43-mCherry, whereas only 35±0.1% of cytoplasmic ΔNLS inclusions recruited TDP-43-mCherry (Figure 5M). However, cytoplasmic ΔNLS/4FL or ΔNLS/2KQ variants exhibited 88.47±2.31% and 77.52±4.49%, respectively, of inclusions sequestering endogenous TDP-43 (Figure 5M), indicating that disease-associated pathological changes increase the sequestration of normal nuclear TDP-43 to cytoplasmic inclusions. Finally, smaller WT puncta or anisosomes formed by 4FL and 2KQ mutants in the nucleus almost always recruited endogenous TDP-43-mCherry (Figure 5N). In contrast, puncta in cells expressing cytoplasmic TDP-43-GFP mutants ΔNLS (28.31±11.54%), ΔNLS/4FL (25.95±1.29%), or ΔNLS/2KQ (18.65±2.82%) showed a significantly lower proportion of TDP-43-mCherry sequestration (Figure 5N).

These results show that nuclear RNA-binding-ablated or acetylation-mimicking mutant TDP-43 proteins form an increased number and size of anisosomes compared to WT TDP-43-GFP puncta. Nuclear puncta or anisosomes formed from pathological TDP-43-GFP sequester endogenous TDP-43 in a concentration-dependent manner, which depletes diffuse nuclear endogenous TDP-43 seen earlier (Figure 5D, F; Supplementa y Figure 5D). Additionally, we found that RNA-binding-ablated or acetylation-mimicking mutants in the cytoplasm led to significantly greater sequestration of endogenous TDP-43, compared to ΔNLS alone. Together these findings suggest that changes in RNA-binding capacity and acetylation of TDP-43 that occurs in ALS and FTD, enhances the propensity for self-associated into puncta and depletes free nuclear TDP-43. Immunostaining for total TDP-43 also revealed that anti-TDP-43 encircled TDP-43-GFP aggregates but was unable to stain the core, while fluorescent signals from endogenous TDP-43-mCherry featured throughout these structures (Supplementary Figure 8), as shown previously (Yu et al., 2021). While anti-TDP-43 antibodies may not penetrate inclusions, this provides evidence that free normal nuclear TDP-43 is not only sequestered to the surface, but likely ‘co-aggregates’ with over-expressed exogenous TDP-43-GFP proteins during inclusion formation. Importantly, this sequestration into large inclusions, puncta, or anisosomes may mark critical molecular events by which normal nuclear TDP-43 becomes mislocalised, dysfunctional, and pathological in cells following initiation of TDP-43 aggregation.

### Free normal nuclear TDP-43 is specifically sequestered into acetylation-mimicking TDP-43 inclusions, becoming insoluble and immobile

As acetylation, along with other PTMs, is a key feature of TDP-43 pathology in ALS and FTD, we next sought to determine whether sequestration of normal TDP-43 to acetyl-mimicking mutant TDP-43 structures altered its solubility or mobility, indicating dysfunction and pathological transition. We performed phospho-TDP-43(403/404) immunostaining on *TARDBP-mCherry* cells expressing nuclear or cytoplasmic acetylation-mimic TDP-43-GFP proteins to further characterise the sequestration of endogenous TDP-43 into puncta and inclusions. Confocal imaging demonstrates that a majority of large nuclear or cytoplasmic acetylation-mimicking mutant TDP-43 inclusions that sequester normal TDP-43-mCherry were highly phosphorylated (Figure 6A,B, white arrowheads). Smaller cytoplasmic puncta formed by acetylated TDP-43 showed less TDP-43-mCherry sequestration and were occasionally phosphorylated (Figure 6B). Nuclear anisosomes that readily recruited free normal nuclear TDP-43 were never phosphorylated (Figure 6A, yellow arrowheads). The tendency for large, phosphorylated acetylation-mimickingTDP-43 inclusions to recruit endogenous TDP-43 prompts investigation to determine whether this sequestration constitutes a pathological transition of normal TDP-43 protein, leading to insolubility and immobility.

**Figure 6.**
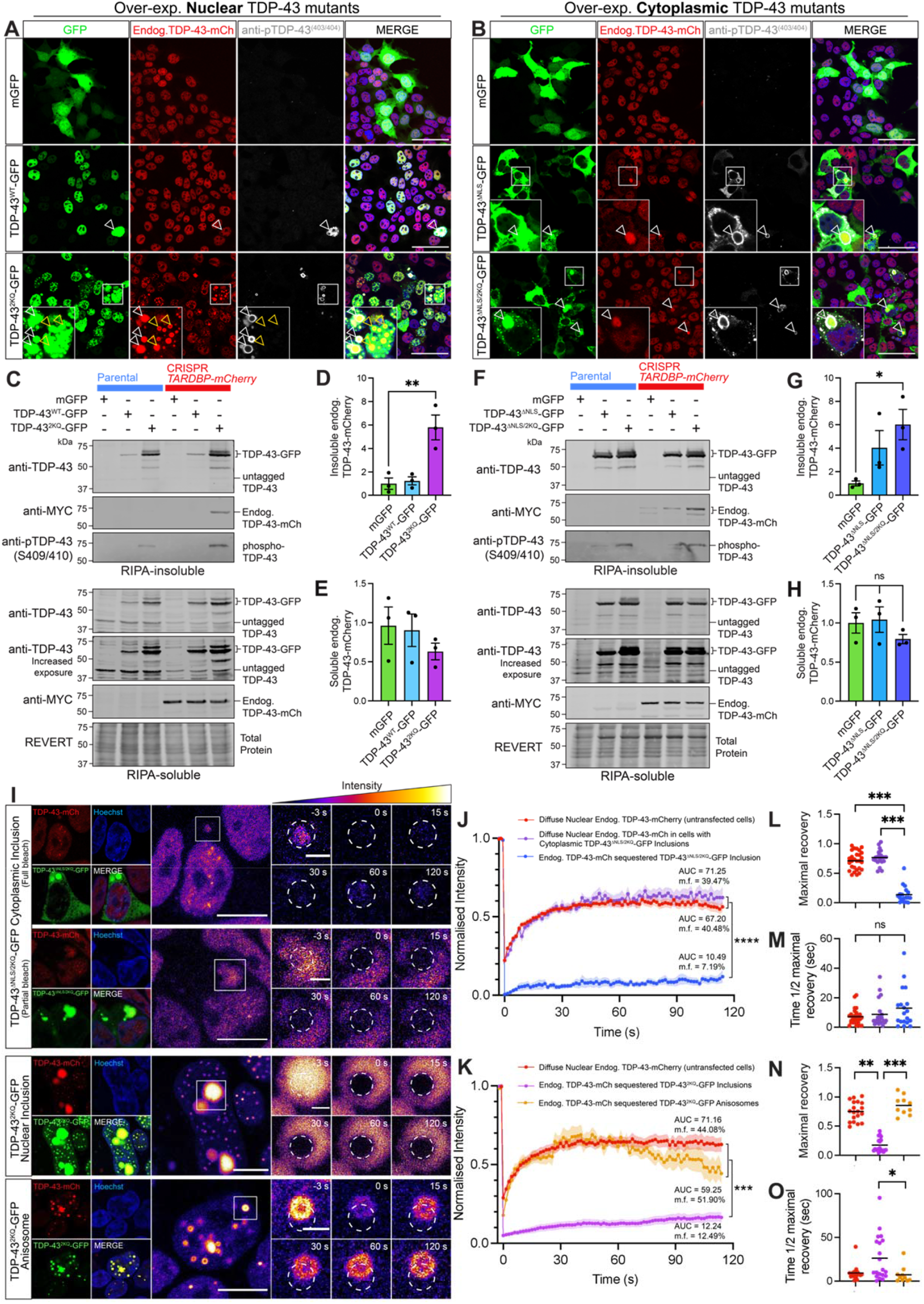
Sequestered endogenous TDP-43 becomes insoluble and immobile in nuclear and cytoplasmic inclusions, however retains high protein mobility when recruited to nuclear anisosomes. **(A)** Representative confocal micrographs of CRISPR *TARDBP-mCherry* cells expressing nuclear TDP-43^WT^-GFP, acetylation-mimic TDP-43^2KQ^-GFP, or mGFP control. **(B)** Confocal micrographs of cytoplasmic TDP-43^ΔNLS^-GFP, acetylation-mimic TDP-43^ΔNLS/2KQ^-GFP, or mGFP control, 48h post-transfection. White arrowheads indicate inclusions comprising endogenous over-expressed and phosphorylated TDP-43. Yellow arrowheads indicate TDP-43 anisosomes which lacked phosphorylated TDP-43. **(C)** Immunoblotting of CRISPR *TARDBP-mCherry* cells expressing nuclear or **(F)** cytoplasmic mutants, demonstrate changes in endogenous TDP-43-mCherry levels within **(D**,**G)** RIPA-insoluble and **(E**,**H)** -soluble protein fractions, respectively, quantified by densitometry analysis. **(I)** FRAP was used to quantify the mobility of endogenous TDP-43-mCherry proteins recruited to distinct nuclear or cytoplasmic structures formed by the over-expression of acetylation-mimic TDP-43-GFP mutants. Scale bars on left-hand panels = 10 μm; zoomed right-hand panels = 2 μm. Pseudocolouring demonstrates changes in endogenous TDP-43-mCherry intensity, which was quantified within the bleaching region-of-interest (white dotted circle) for 120s post-bleaching. FRAP analysis of the normalised intensity during the recovery period for cells expressing **(J)** cytoplasmic or **(K)** nuclear acetylation-mimic mutant TDP-43-GFP, along with mobility fraction (m.f.) values and area-under-the-curve (AUC) averaged for each cell, which was used for statistical comparison. **(L**,**N)** Maximal fluorescence intensity observed at the final time-point (120s), and **(M**,**O)** time taken to recover to half of the maximal fluorescence level for nuclear and cytoplasmic TDP-43 mutants respectively. Same colour scheme as the intensity line graphs, with each point representing a single cell value, and the black line representing the mean across *n*=3 biologically independent repeats. Statistically significant differences between the means were analysed using one-way ANOVA multiple comparisons with Tukey’s post hoc test (* = p < 0.05; ** = p < 0.01).

We therefore analysed the levels of detergent-soluble and -insoluble levels of normal endogenous TDP-43 in cells over-expressing these pathological TDP-43-GFP mutants. Over-expression of either nuclear or cytoplasmic acetylation-mimic TDP-43-GFP (Figure 6C,F) resulted in a significant 6-fold increase of insoluble endogenous TDP-43 levels, detected via the the myc tag incorporated during CRISPR editing (Figure 6D,G). Soluble endogenous TDP-43-mCherry levels were not significantly altered (Figure 6E,H). These findings show that nuclear and cytoplasmic phosphorylated inclusions formed by acetylation-mimicking TDP-43 sequester, and render insoluble, normal TDP-43.

We next investigated whether this sequestration alters the mobility dynamics of normal TDP-43-mCherry (Figure 6I). As expected, FRAP analysis revealed that the incorporation of endogenous TDP-43 into acetylated cytoplasmic TDP-43 inclusions abolishes its mobility, in comparison to diffusely distributed nuclear TDP-43 (Figure 6J). Either complete or partial bleaching of endogenous TDP-43 within nuclear or cytoplasmic inclusions both showed minimal fluorescence recovery, indicating almost no dynamic exchange with surrounding cytoplasmic proteins, or within the inclusion itself, respectively. Specifically, endogenous TDP-43 sequestered into large cytoplasmic TDP-43^ΔNLS/2KQ^-GFP inclusions showed an AUC of 10.49 a.u. and m.f. of 7.19%, a maximal recovery of 12.63% (Figure 6L) and time to reach half maximal (t1/2) of 17.17 s (Figure 6M). In some cells, acetyl-mimicking TDP-43-GFP in the nucleus formed large inclusions that recruited endogenous TDP-43-mCherry, which also showed severe mobility impairment (Figure 6K), demonstrated by AUC of 12.24 a.u., m.f. of 12.49, and significant decreases in maximal recovery of 14.70% (Figure 6N), and t1/2 of 26.18 s (Figure 6O). In contrast, endogenous TDP-43-mCherry incorporated more frequently into nuclear TDP-43^2KQ^-mGFP anisosomes and retained high mobility and rapid rate of recruitment (Figure 6K), with AUC of 59.25 a.u., m.f. of 51.90%, maximal recovery of 37% (Figure 6L) and t1/2 of 6.30 s (Figure 6M). Dynamic properties of endogenous TDP-43 in anisosomes did not greatly differ overall from diffusely distributed endogenous TDP-43 in untransfected cells. This was likely due to the high concentration of nuclear acetylation-mimicking TDP-43 which formed anisosomes that readily recruited endogenous TDP-43. We note that the variability through time and decline following peak fluorescence recovery for acetylated TDP-43 anisosomes (Figure 6K) is likely due to movement of these structures within the nucleus partially out of the FRAP region-of-interest. Together, these findings suggest that endogenous TDP-43 is specifically sequestered into a variety of pathological TDP-43 structures to significantly alter the solubility and mobility dynamics of endogenous TDP-43. Endogenous TDP-43 incorporated into large, phosphorylated nuclear or cytoplasmic acetyl-mimicking TDP-43 inclusions correlated strongly with decreased solubility, along with a dramatic loss of mobility, while endogenous TDP-43 within anisosomes comprised of acetyl-mimicking TDP-43 remained highly mobile.

## Discussion

It has remained unclear whether the accumulation of pathological TDP-43 proteins in ALS and FTD can directly divert normal TDP-43 away from its functional pool, to contribute to TDP-43 nuclear depletion and loss-of-function. Here, we developed novel cellular tools to study changes in normal nuclear TDP-43 abundance, localisation, self-assembly, inclusion formation, and mobility dynamics, under disease-mimicking conditions in real-time. Notably, we found that introduction of RNA-binding-deficient or acetyl-mimicking TDP-43 in either the nucleus or cytoplasm progressively decreases free nuclear TDP-43 abundance in a concentration-dependent manner. Endogenous TDP-43 was specifically sequestered into large cytoplasmic or nuclear inclusions formed by acetylation-mimicking TDP-43 variants, becoming insoluble and immobile. However, nuclear acetylation-mimicking TDP-43 also spontaneously formed TDP-43 anisosomes (Yu et al., 2021), which readily sequestered over half of total endogenous TDP-43 levels, but remained soluble and mobile. Endogenous wild-type TDP-43 also formed diverse highly dynamic biomolecular condensates via LLPS under various conditions. Indeed, arsenite-mediated oxidative stress increased the formation of nuclear liquid-like droplet structures with similar mobility dynamics to physiological puncta observed in unstressed cells. We also found that oxidative stress also induced LLPS formation of endogenous TDP-43 anisosomes in the nucleus, which notably were not formed by over-expressed wild-type TDP-43. These findings indicate that RNA-binding-deficient and acetylated TDP-43 proteins, or disease-associated oxidative stress, drive recruitment of normal nuclear TDP-43 to insoluble inclusions or other condensed assemblies in a compartment-specific manner. This mechanism of pathological sequestration likely decreases the availability of functional nuclear TDP-43, which may critically exacerbate TDP-43 dysfunction in ALS and FTD.

A loss of free nuclear TDP-43 in disease may be triggered by cellular stressors that induce nuclear-to-cytoplasmic redistribution or phase transitions of endogenous TDP-43 into condensed assemblies. Phase-separated TDP-43 proteins comprising stress-induced liquid droplets, SGs, and anisosomes likely remain soluble and dynamic, however confinement within these reversible structures may alter physiological TDP-43-RNA or -protein interactions (Lee et al., 2022; Yu et al., 2021). Furthermore, chronic induction, abnormal assembly, or aberrant phase transitions of TDP-43 condensates may seed intranuclear or cytoplasmic aggregation, as observed in degenerating neurons (Chen and Cohen, 2019; Gasset-Rosa et al., 2019; Li et al., 2013; Mann et al., 2019; Yu et al., 2021; Zhang et al., 2019). Critically, TDP-43 LLPS has largely been studied by over-expressing TDP-43 constructs with mutations, truncations, or optogenetic clustering tags that promote self-assembly. Alternatively, in this study we sought to characterise these LLPS-related structures formed by endogenous TDP-43, to more closely study the physiological context. We consistently observed distinct liquid-like ‘nuclear bodies’ or ‘droplets’ under physiological conditions, but found that arsenite-mediated oxidative stress increased their formation, and also induced cytoplasmic G3BP1-positive TDP-43 SGs, aligning with previous studies of exogenous TDP-43 LLPS (Gu et al., 2021; Wang et al., 2020). Notably, we found that oxidative stress increased the formation of rare endogenous TDP-43 anisosomes, of which spontaneous formation has been previously reported upon over-expression of RNA-binding-ablated or acetylation-mimicking TDP-43 variants (Yu et al., 2021). Combined treatment with histone deacetylase and proteasome inhibition has also been shown to induce endogenous TDP-43 anisosome formation in rats (Yu et al., 2021), suggesting that diverse cellular stressors can stimulate TDP-43-containing anisosome formation. We found that endogenous TDP-43 anisosomes form rapidly and have high mobility in live cells, and that TDP-43 anisosome formation can be induced by oxidative stress, without requiring RNA-binding mutations or proteolytic stress. We speculate that cellular stress alters TDP-43-RNA interactions, leading to LLPS and dynamic self-assembly into RNA-deficient anisosomes, to deplete the free nuclear pool of functional TDP-43 (Cohen et al., 2012; Markmiller et al., 2021). It remains unclear whether other cellular stressors trigger endogenous TDP-43 anisosome formation, and whether the assembly of RNA-binding-competent TDP-43 into anisosomes impacts RNA-regulatory functions.

Accumulation of pathological TDP-43 itself likely drives nuclear TDP-43 depletion by decreasing endogenous TDP-43 expression via auto-regulation (Ayala et al., 2011b; Hallegger et al., 2021; Koehler et al., 2022), or inducing mislocalisation by disrupting processes such as nucleocytoplasmic transport (Chou et al., 2018). However, potentially independent of these mechanisms, we found striking changes in endogenous TDP-43 distribution within cells with high abundance and aggregation of exogenous TDP-43-GFP. Notably, endogenous TDP-43-mCherry was sequestered into inclusions or puncta formed by pathological TDP-43-GFP variants, suggesting that a self-perpetuating cycle of pathological conversion may drive TDP-43 dysfunction in disease separately from or in addition to RNA-mediated auto-regulatory perturbations. Indeed, single-cell analysis revealed that nuclear RNA-binding-ablated (4FL) or acetylation-mimicking (2KQ) variants in particular sequestered a large proportion of endogenous TDP-43 into anisosomes, in a concentration-dependent manner. These findings indicate that as pathological TDP-43 accumulates over time, more and more normal TDP-43 may be diverted from normal nuclear function. In the cytoplasm, we similar found that the addition of pathology-mimicking 4FL or 2KQ mutations increased inclusion and puncta formation over and above the ΔNLS-only variant, however with only occasional recruitment of endogenous TDP-43, indicating compartment-specific tendencies for sequestration. While over-expression of TDP-43-GFP variants did not significantly alter total endogenous TDP-43 levels in this cell system, we showed that the dramatic redistribution into mutant inclusions, anisosomes, or puncta completely accounted for the loss of diffuse nuclear endogenous TDP-43. Our study therefore highlights that concentration-dependent sequestration of normal TDP-43 to pathological structures is likely a crucial concurrent process driving nuclear depletion of TDP-43 in ALS and FTD. Such sequestration, or ‘co-aggregation’, of over-expressed WT TDP-43 has been demonstrated upon introduction of disordered or mislocalised TDP-43 proteins, with scarce evidence of endogenous human TDP-43 being sequestered in cell lines (Che et al., 2015; Gasset-Rosa et al., 2019; Winton et al., 2008; Zhang et al., 2013). Crucially, our unbiased quantitative image analysis demonstrated direct correlations between mutant TDP-43 aggregation and the sequestration and depletion of free normal nuclear endogenous TDP-43, which was amplified by acetylated or RNA-binding-ablated TDP-43 mutants. While previous studies indicate that loss of RNA-binding promotes aberrant TDP-43 self-assembly and aggregation (Pérez-Berlanga et al., 2022), others suggest that TDP-43-RNA interactions mediate such sequestration, as RNA-binding ablated 4FL mutants were shown to not co-precipitate with endogenous TDP-43 (Che et al., 2015; Jiang et al., 2022). However, our results indicate that RNA-binding function is not necessary for sequestration of normal TDP-43. It is unclear from our study whether nuclear acetylation-mimicking TDP-43 retains RNA-binding capacity, or whether nuclear RNA abundance influences the formation of these TDP-43 assemblies. Therefore, the role of RNA-binding capacity in aggregation and endogenous TDP-43 sequestration requires further investigation, with additional mechanisms likely involved.

Since we found that pathological TDP-43 could recruit normal nuclear TDP-43 to large inclusions, puncta or anisosomes in live cells, we also sought to determine whether this sequestration caused a transition to a pathological state. Inclusions formed by over-expressed acetyl-mimic TDP-43 that contained endogenous TDP-43 were often phosphorylated, reminiscent of end-stage disease pathology (Brettschneider et al., 2013), which may be promoted by acetylation (Wang et al., 2017). Furthermore, expression of acetylation-mimicking TDP-43 increased the levels of insoluble endogenous TDP-43 and phosphorylated TDP-43, concurrent with a slight decrease in soluble endogenous TDP-43. Importantly, sequestration of normal TDP-43 to nuclear or cytoplasmic inclusions led to insolubility and loss of mobility, likely indicating irreversible pathological transitions and loss-of-function, despite no change in total endogenous TDP-43 levels. Previous work has demonstrated co-precipitation of endogenous TDP-43 with mutant TDP-43 C-terminal fragments (Che et al., 2015), with others showing that over-expression of cytoplasmic ΔNLS mutant TDP-43 increases insoluble levels of endogenous TDP-43 (Winton et al., 2008). We are first to report sequestration of endogenous normal nuclear TDP-43 to various RNA-deficient inclusions, anisosomes, or puncta in either the nucleus or cytoplasm of live cells, and show that this recruitment alters dynamic properties of endogenous TDP-43 to conform to exogenous mutant TDP-43 proteins that rapidly form these structures.

While more dramatic sequestration was caused by acetylated TDP-43 anisosomes, the pathophysiological implications remain unclear. Normal TDP-43 within anisosomes retained liquid-like properties and dynamic exchange with the free nuclear pool of TDP-43, corresponding with high mobility of over-expressed TDP-43^2KQ^-mClover anisosomes previously characterised (Yu et al., 2021). However, it is unknown whether normal TDP-43 recruited to anisosomes can interact with RNA, whether acetylated TDP-43 anisosomes inhibit or seed TDP-43 aggregation, and whether anisosomes form in human ALS and FTD (Yu et al., 2021). In either case, it is likely that sequestration constitutes a significant disruption of normal nuclear TDP-43 localisation, availability, and physiological interactions, contribute significantly to loss-of-function mechanisms of neurodegeneration, independently of changes to overall levels of normal nuclear TDP-43. Combined, these findings indicate that TDP-43 acetylation not only increases aggregation propensity, but also drives sequestration and pathological transitions of normal TDP-43, to potentiate disease-associated TDP-43 nuclear depletion, dysfunction, and pathology formation in ALS and FTD.

Studying endogenous TDP-43 protein over time is advantageous for elucidating disease-relevant mechanisms driving depletion and aggregation of the native TDP-43 protein, which becomes pathological in almost all cases of ALS and half of FTD (Ling et al., 2013). Our CRISPR-tagging approach allows for live-cell visualisation of endogenous TDP-43 in real-time, overcoming limitations of over-expression studies, end-point analyses and traditional staining techniques, to reveal assembly and composition of complex TDP-43 structures. CRISPR/Cas9-mediated fluorescent tagging of endogenous *TARDBP* has been reported in HEK293T cells for live-cell imaging (Jiang et al., 2022), in human neuroblastoma cells for FRAP (Gasset-Rosa et al., 2019) or in iPSCs for real-time quantification of nuclear endogenous TDP-43 (Chua et al., 2022; Gasset-Rosa et al., 2019; Jiang et al., 2022). Our unique CRISPR method, whereby one multiplexed editing reaction mediated insertion of two different fluorescent tags across different *TARDBP* alleles, demonstrates potential for differential fluorescence tagging of multiple distinct endogenous proteins for concurrent intracellular analysis. Recently, CRISPR-tagged endogenous myc-TDP-43 zebrafish models have been developed (Petel Légaré et al., 2022), supporting feasibility of fluorescently-tagged endogenous TDP-43 *in vivo* models. To our knowledge, this is the first endogenous-tagging application of mAvicFP1 (Lambert et al., 2020), which favourably exhibits high fluorescence intensity and photostability, and is amenable to single-molecule super-resolution microscopy (Gavrikov et al., 2020). Future directions in the use of our *TARDBP-mAvicFP1/-mCherry* cell models include CRISPR-screening applications, live-cell single-molecule imaging or specific mutations of endogenous TDP-43. Finally, as we characterised dynamic TDP-43 structures (e.g. anisosomes) that were not formed by over-expressed TDP-43, it will be interesting to see whether novel dynamic TDP-43 assemblies could be discovered with high-resolution real-time imaging.

## Conclusions

To interrogate molecular mechanisms by which TDP-43 protein becomes mislocalised and aggregated in disease, this study sought to investigate whether disease-associated stressors or pathology-mimicking TDP-43 variants alter endogenous TDP-43 abundance, localisation, self-assembly, aggregation, and mobility dynamics over time. We found that oxidative stress induces dynamic self-assembly of endogenous TDP-43 into ‘anisosomes’. Notably, we demonstrated that accumulation and aggregation of RNA-binding-ablated or acetylation-mimicking TDP-43 drives nuclear depletion of normal TDP-43, via concentration-dependent sequestration into inclusions, puncta or anisosomes. This specific sequestration into large, phosphorylated nuclear or cytoplasmic inclusions formed by pathological TDP-43 led to insolubility and immobility of normal TDP-43, indicating irreversible pathological transition. The most dramatic sequestration was caused through formation of nuclear anisosomes comprised of acetylation-mimicking TDP-43, although, interestingly, endogenous TDP-43 recruited to these dynamic structures remained highly mobile. Our results suggest that changes in localisation and availability of TDP-43 via sequestration into either disease-reminiscent inclusions or dynamic phase-separated structures may drive TDP-43 dysfunction and perpetuate TDP-43 aggregation in ALS and FTD.

## Materials and Methods

Chemicals were sourced from Sigma unless otherwise stated.

### CRISPR/Cas9-mediated fluorescent protein knock-in to endogenous TARDBP genomic locus

CRISPR/Cas9 gene-editing was used to generate endogenous TDP-43-tagged HEK293T cell lines with the insertion of fluorescent proteins, mAvicFP1 and mCherry, at the *TARDBP* C-terminus. The coding sequence for mAvicFP1, a monomeric, bright and photostable green fluorescent protein (Lambert et al., 2020), or the red fluorescent protein mCherry, were integrated immediately upstream of the exon 6 STOP codon, with a dual DYK*-* and Myc-tag linker sequence (Figure 1A-D). Gene insertion was mediated by homologous recombination of the PCR amplicon DNA donor template, which contained the fluorescent tag insertion sequence flanked by proximal and recessed homology arms of 50 and 56bp, respectively.

### Single guide RNA and donor template primer design

sgRNA oligonucleotides and dsDNA donor templates for mAvicFP1/mCherry insertion were designed using the program *Benchling* and synthesised by *GenScript*. The sgRNA sequence (5’-GCTGGGGAATGTAGACAGTG-3’) was designed to anneal to the *TARDBP* C-terminus and direct Cas9-mediated double-stranded break (DSB) 3 base-pairs (bp) upstream of the PAM site. Donor templates featured the insertion sequences of *mAvicFP1* or *mCherry*, preceded by a mutation of the PAM site (GGG>GGA) to prevent future Cas9-DSBs after successful gene-editing, and followed by FLAG-myc linkers to separate fluorescent protein tags from TDP-43 to ensure normal folding. Insertion sequences were flanked by 50 bp recessed or 56 bp proximal homology arms (Has) that were complementary to the endogenous *TARDBP* C-terminal sequence, to specifically direct insertion 6 bp upstream of the DSB. The donor templates were designed such that *FLAG* and *myc* sequences comprised the linker sequence between *TARDBP* and the fluorescent protein for downstream analysis of the edited protein with FLAG or myc antibodies.

### PCR amplification of donor DNA templates for homology-directed repair

Donor templates for homology-directed repair were produced as linear DNA fragments from PCR amplification, as supplying PCR amplicons of just the insertion sequence produces greater editing efficiency compared to circular DNA constructs, and generates a higher concentration of DNA template (Paix et al., 2017). Donor template PCR products were purified using the QIAquick PCR Purification kit (QIAGEN, 28104), eluted in 15 μL to maximise DNA concentration. The expected DNA band size of these amplicons (mAvicFP1 877 bp, mCherry 871 bp) was confirmed by analysing dsDNA donor templates on a 1% (w/v) agarose gel at 85 V for 1 h.

### RNP complex assembly & electroporation delivery

Functional sgRNAs were formed by hybridisation of crRNA and tracrRNA (5’-AGCAUAGCAAGUUAAAAUAAGGCUAGUCCGUUAUCAACUUGAAAAAGUGGCACCGAGUCGGUGCUUU-3’), whereby lyophilised crRNA and tracrRNA oligonucleotides were each reconstituted to 100 μM in sterile Tris-EDTA, and then combined in equimolar ratio to a final concentration of 40 μM, diluted with duplex buffer (100 mM Potassium Acetate, 30 mM HEPES, pH 7.5). This duplex was heated at 95°C for 5 min and cooled at room temperature for 2 h, then diluted to 10 μM in Tris-EDTA. RNP complexes were created by mixing 1.625 μL cr:tracrRNA duplex (10 μM) with 2.6 μL purified NLS-Cas9-NLS protein (6.25 μM) gently by pipetting, and incubating for 10-60 min.

300,000 HEK293 cells were aliquoted for each electroporation condition (DNA-free negative control, *mAvicFP1/mCherry* PCR-amplicon donor templates and pEGFP-N1 positive control). The RNP complex and donor DNA was delivered to the cells by electroporation using the Neon Transfection System 100 μL kit (ThermoFisher, MPK10025) with Electrolytic-Buffer-E2. Cells were resuspended in Buffer-R and transferred to the corresponding RNP complex tubes. 2.6 μg of each dsDNA donor template was used. HEK293 cells/RNP complexes/donor templates were mixed and aspirated into a 100 μL Neon tip and electroporated with 1200 V, 2 pulses, 30 ms pulse width. The pipette was immediately removed from the device and cells transferred to a 12-well plate prepared with 1 mL warm media, and incubated for 3 days.

### FACS isolation of successfully edited cells and selection of monoclonal cell lines

CRISPR/Cas9-edited cells were sorted using the BD Influx FACS with 488 nm and 561 nm lasers for excitation of mAvicFP1 and mCherry, respectively. Quandrant gating defined cell populations that were mAvicFP1-positive, mCherry-positive, and ‘dual-tagged’ mAvicFP1-positive/mCherry-positive, which were seeded as single cells into 96-well plates. Single cells were allowed to grow into monoclonal colonies and scanned for mAvicFP1 and mCherry fluorescence intensity using high-content microscopy, which distinguished low-, medium-, and high-intensity TDP-43-tagged cell lines. High-intensity cell lines *TARDBP-mAvicFP1(H8), TARDBP-mCherry(B8)*, and *TARDBP-mAv/-mCh(C4)* were utilised for subsequent experiments throughout this study.

### Sanger sequencing

Genomic DNA was extracted from approximately 1,000,000 cells of the CRISPR-edited *TARDBP-mAvicFP1(H8), TARDBP-mCherry(B8)*, or *TARDBP-mAv/-mCh(C4)* HEK293 lines with a Bioline Isolate II Genomic DNA Kit [BIO-52066]. For each cell line, a ∼900 bp ‘genotyping zone’, comprising the C-terminal *TARDBP* fluorescent tag insertion site was amplified by PCR, with specific primers annealing to the upstream *TARDBP* exon 6 sequence and *mAvicFP1* or *mCherry* sequence. PCR products were analysed by DNA electrophoresis on a 1% agarose gel, before gel purification of the expected genotyping template bands, using a Bioline Isolate II Gel & PCR Kit [BIO52059]. Forward or reverse sequencing primers were each combined with 75ng of gel-purified genomic DNA genotyping sequences, and supplied to Australian Genome Research Facility for Sanger sequencing.

### Cell culture and transfection

Human Embryonic Kidney (HEK293T/17; ATCC^®^-CRL-11268) cells were maintained in Dulbecco’s Modified Eagle Medium/Ham F12 media (DMEM/F12) with 10% (v/v) foetal calf-serum (Gibco). Cells were incubated at 37°C, 5% CO_2_ and passaged at 80% confluency, and tested regularly for mycoplasma contamination. Cells were transfected using Lipofectamine 2000 (Invitrogen) and plasmid DNA according to the manufacturer’s instructions. Plasmids included human pLenti-C-mGFP, pLenti-TDP-43^WT^-C-mGFP (Origene), and pLenti-TDP-43^4FL^-C-mGFP, pLenti-TDP-43^2KQ^-C-mGFP, pLenti-TDP-43^ΔNLS^-C-mGFP, pLenti-TDP-43^ΔNLS/4FL^-C-mGFP, or pLenti-TDP-43^ΔNLS/2KQL^-C-mGFP (Origene, mutations by GenScript). Cells were seeded onto coverslips in 24-well plates at a density of 25,000 cells/well and incubated for 24 h. For transfections, 0.5 μg DNA and 0.75 μL Lipofectamine 2000 were used in a total of 100 μL OptiMEM (Gibco) per well. Cells were incubated for 48 h to allow protein expression from the plasmid DNA before analysis by immunoblot or immunocytochemistry.

### Immunoblotting

At experiment end-points, cells were harvested and washed in ice-cold PBS, before soluble and insoluble protein fractions were extracted by sequential RIPA-buffered (50 mM Tris [pH 7.5], 150mM NaCl, 1% [v/v] NP-40, 0.5% [w/v] sodium deoxycholate, 0.1% [w/v] SDS, 1 mM EDTA, 1 mM PMSF, 1 × protease/phosphatase inhibitor cocktail) and urea-buffered protein lysis, Precellys homogenisation and ultracentrifugation. RIPA-soluble protein extracts were quantified using a BCA Protein Quantification Kit (Pierce) and plates were analysed on a ClarioStar Optima plate reader (BMG Lab Technologies). 50 μg of RIPA-soluble protein and an equivalent volume of the corresponding urea-soluble fraction were analyzed by 12 % SDS-PAGE, run for 150 V for 60 min and then transferred to a nitrocellulose membrane (Li-Cor) at 100 V for 90 min. Total protein was quantified with Revert 700 total protein stain [#912-11011, Li-Cor] according to the manufacturer’s instructions, before membranes were blocked in 5% (w/v) BSA Tris buffered saline (0.5 M Tris-base, 1.5 M NaCl) with 0.05% (v/v) Tween-20 (TBS-T). Blots were immunolabelled overnight at 4ºC with primary antibodies, including a rabbit anti-TDP-43 polyclonal antibody (PAb) [1:2000, #10782-2-AP, ProteinTech], mouse anti-myc monoclonal antibody (MAb) [1:500, #2276S (Clone 9B11), Cell Signaling Technologies]. Secondary antibodies included goat anti-mouse IRDye-800 or goat anti-rabbit IRDye-680 conjugated secondary PAbs (Li-Cor). Signals were detected by imaging with the Li-Cor Odyssey Clx Infrared Imaging System, and quantified using Li-Cor Image Studio version 3.1.

### Live-cell imaging

In all live-cell imaging experiments, cell nuclei were stained with Hoechst [1:10000, #33342, Thermo Fisher] for 5 min and washed with fresh media prior to imaging. For 48 h time-course imaging of *TARDBP-mCherry(B8)* cells over-expressing TDP-43-GFP mutants, cells seeded at 30,000 cells/well in 24-well plates were transferred to the Celena-X live-cell imaging microscope (Logos Biosystems), within an insulated chamber at 37°C and 5 % CO_2_. mCherry and GFP channels were imaged for 20 ROIs, every 2h for 48h, at 20X magnification with laser auto-focus in every ROI. Image processing was completed with FIJI and image analyses were conducted using CellProfiler version 4.2.1 (McQuin et al., 2018) using custom pipelines. For 1 h continuous imaging of *TARDBP-mAvicFP1(H8)* cells treated with or without sodium arsenite, 20,000 cells/dish seeded in Cellvis glass-bottom 20 mm dishes [#D29-20-1.5-N] were transferred to the Zeiss LSM 710 confocal microscope and identified with a Plan-Apochromat 63x oil objective (1.4 NA, 190μm WD, 0.19 μm/pixel), blue laser diode for DAPI (405 nm), argon laser for AlexaFluor488 (488 nm), and 561 nm laser for AlexaFluor555 excitation. During imaging, cells were contained within an insulated chamber at 37°C and 5 % CO_2_. The GFP channel was imaged every 1 min for 1 h, at 63X magnification, with Z-stacks of 6-8 slices separated by the recommended optimal distance. Image processing was completed with FIJI, including maximum Z-projections.

### Fluorescence recovery after photobleaching (FRAP)

For FRAP imaging of *TARDBP-mAvicFP1(H8)* cells treated with or without sodium arsenite or *TARDBP-mCherry(B8)* cells over-expressing TDP-43-GFP mutants, cells seeded at 20,000 cells/dish in Cellvis glass-bottom 20 mm dishes [#D29-20-1.5-N] were transferred to the Zeiss LSM 710 confocal microscope, were cells were contained within an insulated chamber at 37°C and 5 % CO_2_. Cells were identified using a Plan-Apochromat 63 × 1.4NA oil objective on the Zeiss LSM 710 confocal microscope, and Zeiss ZEN software was used for data acquisition. FRAP acquisition was performed on cells within a field of view including adjacent cells of similar intensity (to control for photo-bleaching). A baseline of 3 images was acquired (using 4 % argon 488 or 561 nm laser intensity) prior to bleaching. The regions of interest on the cells were bleached for about 10 seconds (200 iterations) using 100 % argon 488 or 561 nm laser intensity over 77 imaging cycles (∼2 minutes), in order to completely deplete fluorescence intensity of endogenous TDP-43 within structures of interest. The image scan speed was set at 10. Data presented are representative of three or more independent experiments. Acquired data was normalised using the web-based resource tool, EasyFRAP (Koulouras et al., 2018). Mobile fractions (m.f.) and recovery time constants (τ) were calculated using Zen software from regions of interest of curve-fitted data.

### Immunofluorescence staining

Cells on glass coverslips were fixed with 4 % paraformaldehyde in phosphate-buffered saline (PBS) for 10 min at room temperature, before blocking & permeabilisation with 3 % BSA in PBS-T (+0.1 % Triton-X100) for 1 h at room temperature. Primary antibodies were diluted in blocking & permeabilisation buffer, including a rabbit anti-TDP-43 PAb [1:1000, #10782-2-AP, ProteinTech], rabbit anti-phospho TDP-43 (ser403/404) PAb [1:1000, CAC-TIP-PTD-P05, CosmoBio], and rabbit anti-G3BP1 PAb [1:500, #13057-2-AP, ProteinTech], and applied overnight at room temperature. Secondary antibodies were similarly prepared, including goat anti-Rabbit H+L AlexaFluor647 [1:500, A-21245, Life Tech], and applied for 1.5 h at room temperature. Cells were counterstained with DAPI [1:1000, #62248, Thermo Fisher] for 10 min at room temperature, mounted onto microscope slides with ProLong Gold Antifade Mountant [#P36934, ThermoFisher], and allowed to cure overnight at room temperature before microscopy.

### Confocal microscopy

Confocal images were captured using either the Zeiss Laser Scanning Microscope (LSM) 510 META or Zeiss LSM 710. Both are equipped with a 63x oil objective (1.4 NA, 190μm WD, 0.19

μm/pixel), blue laser diode for DAPI (405 nm), argon laser for AlexaFluor488 (488 nm), 561 nm laser for AlexaFluor555, and HeNe 633 nm laser for AlexaFluor647 excitation. Images were acquired using Zen software and all acquisition parameters were applied consistently for all samples in an experimental set. Image processing and intensity profile analyses were completed with FIJI.

### Statistical analysis

Data were represented as mean standard error about the mean (SEM). GraphPad Prism 9 was used for statistical analysis and preparation of graphs, with p-values < 0.05 deemed statistically significant. Live-cell imaging of nuclear TDP-43-mCherry expression over time included n=3 independent repeats, recording 12 ROIs at 20X magnification for each experimental condition. FRAP experiments also included n=3 independent repeats, with at least 5 cells per replicate for each of the distinct TDP-43 assemblies studied. All immunoblot experiments included n=3 independent repeats. Individual data points for each independent replicate are shown overlaid with a line at the mean. Statistically significant differences between the means were determined using a two-tailed paired t-test (where indicated in the figure legend) for comparisons between only two groups, or a one-way ANOVA with a Tukey’s post hoc test for multiple comparisons in experiments with several groups.

## Acknowledgements

This work was supported by the National Health and Medical Research Council (RD Wright Career Development Fellowship 1140386 to AKW), the Ross Maclean Fellowship, and the Brazil Family Program for Neurology. SK was supported by a Thornton Foundation PhD Scholarship and a MND Research Australia PhD Scholarship top-up grant. ATB was supported a Race Against Dementia– Dementia Australia Research Fellowship. RSG was supported by a FightMND Early Career Research Fellowship. Imaging was performed at the Queensland Brain Institute’s Advanced Microscopy Facility, generously supported by the Australian Government through the ARC LIEF grant LE130100078. Fluorescence-activated cell sorting was performed at the Queensland Brain Institute’s Flow Cytometry Facility. We thank Rowan Tweedale and members of the Neurodegeneration Pathobiology Lab for critical evaluation of the manuscript.

## Author Contributions

SK, RSG, and AKW conceived the research; SK, ATB, RSG, and AKW designed the project. SK, ATB and RSG performed the experiments; SK and ATB analysed the data and prepared the figures; SK, RSG, and AKW interpreted the results and wrote the manuscript. All authors read and approved the final manuscript.

